# Crosslink: A fast, scriptable genetic mapper for outcrossing species

**DOI:** 10.1101/135277

**Authors:** Robert J. Vickerstaff, Richard J. Harrison

## Abstract

**Summary:** Crosslink is genetic mapping software for outcrossing species designed to run efficiently on large datasets by combining the best from existing tools with novel approaches. Tests show it runs much faster than several comparable programs whilst retaining a similar accuracy.

**Availability and implementation:** Available under the GNU General Public License version 2 from https://github.com/eastmallingresearch/crosslink

**Supplementary information:** Supplementary data are available at *Bioinformatics* online and from https://github.com/eastmallingresearch/crosslink/releases/tag/v0.5.

## 1 Introduction

Genetic maps are valuable for such key tasks as quantitative trait loci discovery (Hancock et al., 2016), map-based cloning (Huang et al., 2003) and genome assembly scaffolding (Fierst, 2015). Modern high-throughput genotyping technologies routinely provide more markers than can be conveniently mapped using older mapping programs designed around smaller datasets. Even with the ongoing advances in long read sequencing technology the final stage of many genome assemblies is likely to rely on a genetic map for some time to come. This motivates continuing efforts to improve mapping algorithms to facilitate analyses of larger datasets and to increase the level of automation and reliability. Here we focus specifically on the case where the map is constructed using an outcross between two heterozygous parents, which requires a more complex mapping approach than traditional inbred-based F2 or backcross mapping (van Ooijen & Jansen, 2013).

We present Crosslink, an outcross mapper which adapts the minimum spanning tree (MST) method from the inbred mapper MSTmap to provide a rapid initial approximate marker ordering. It also modifies the Gibbs sampler method from outcross mapper JoinMap 4.1 (van Ooijen, 2011) to propagate information unidirectionally leading to shorter convergence times for imputing the missing information inherent in outcross markers heterozygous in both parents. Time complexity of the genetic algorithm used to finalise the marker order is improved by adopting differential recombination counting, such that only the markers adjacent to the break points created by the reordering mutations need to be examined to decide whether to accept or reject the mutation instead of rescanning the entire ordering. Calculation of recombination counts between marker pairs is expedited by converting genotype calls to bit strings with masks representing missing information, counts are then stored in a cache to prevent redundant recalculation for previously examined marker pairs.

## 2 Methods

Crosslink consists of two main programs: crosslink_group which performs marker phasing, missing genotype call imputation using the k nearest-neighbour method (Troyanskaya et al., 2001), formation of linkage groups and approximate marker ordering; and crosslink_map which performs final marker ordering and Gibbs sampler imputation of missing information to allow multi-point recombination fractions to be calculated. Further details are given in the supplement section 1. Input files are encoded using the genotype code conventions of JoinMap (see page 2 of the manual https://github/eastmallingresearch/crosslink/blob/docs/crosslink_manual.pdf and the sample_data directory of the github repository).

## 3 Results

Multiple simulated diploid outcross data sets were generated, each for a single 100 centimorgan chromosome and 200 progeny, varying either error/missing data rates or marker density (see supplement and Fig.1 for details) and used to compare Crosslink to LepMap version Lep-MAP2 v0.2 (Rastas, Paulin, Hanski, Lehtonen, & Auvinen, 2013), OneMap version onemap_2.0-4 with R version 3.1.2 stable version (Margarido, Souza, & Garcia, 2007), Tmap version 1.1 (Cartwright, Troggio, Velasco, & Gutin, 2007) and MSTmap (all tests ran using automated scripts on a high performance Linux computer cluster). LepMap and Tmap used their default settings, OneMap used unidirectional growth mode. Crosslink was tested using either approximate ordering only (running crosslink_group but not crosslink_map) or full ordering (cross-link_group then crosslink_map). MSTmap was tested by prephasing and recoding the data into separate maternal and paternal backcross-type inputs, and then combining the two output maps into a consensus. These extra steps were not counted as part of the execution time. n=8 matched replicates were used for all treatments except for LepMap, Tmap and OneMap at 1000 markers (n=3) or above (n=0) due to excessive run times. JoinMap 4.1 maximum likelihood mapping was also tested, running manually under Windows XP on a desktop computer (therefore running times are not strictly comparable) testing only the first replicate (n=1) for all treatments up to 5000 markers (above which memory was exhausted). Ordering error was measured as one minus the correlation coefficient between the true and reconstructed marker positions. See Fig. 1 for the results.

**Fig 1.**
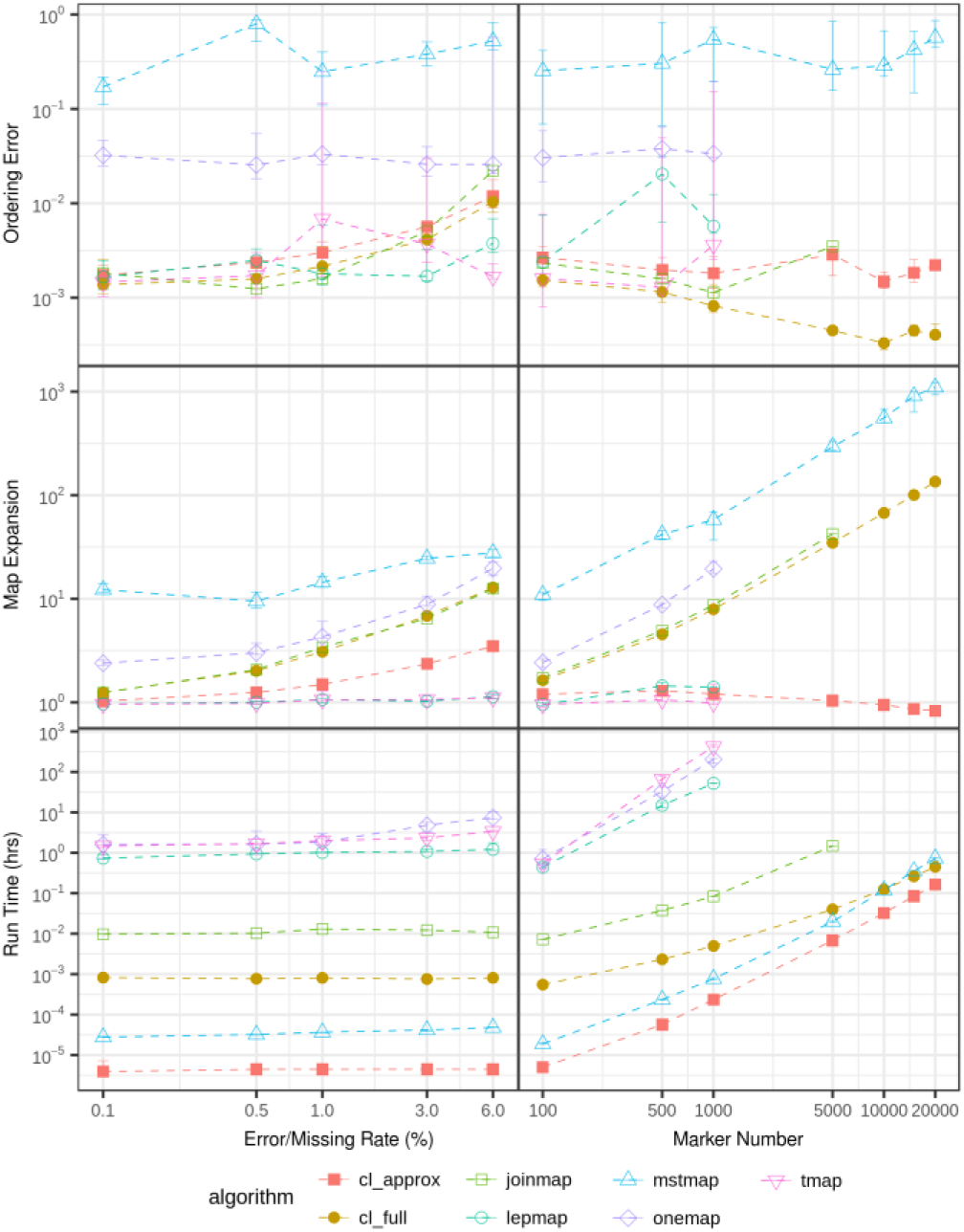
Ordering error, map expansion and running time of Crosslink tested using approximate and full ordering modes, compared to five other mapping programs. Performance is plotted while varying either the genotype-call error and missing rates (left column) or the marker density (right column). Shown are median values of up to n=8 matched replicates plus/minus one quartile. Raw data were from simulated diploid outcrosses with 200 progeny, one 100 cM linkage group, 28% shared, 36% maternal, 36% paternal markers. Error rate experiment: 150 markers (1.5/cM); marker density experiment: error rate 0.5%, missing rate 0.7%.

## 4 Conclusion

Crosslink (in common with JoinMap) was more sensitive to errors and missing data than LepMap and Tmap (which include their own error correction systems), but demonstrated comparable accuracy at low error rates, ran significantly faster and scaled more efficiently than any other program tested including MSTmap. Future development could elucidate whether incorporation of error correction can improve the robustness without significantly increasing run time.

Crosslink was originally developed to map microarray data from cultivated strawberry (*Fragaria* x *ananassa*). We encountered some classes of genotyping errors which we interpreted to be a consequence of the allo-polyploid nature of the strawberry and which we resolved with several additional filtering steps incorporated into Crosslink, otherwise the data were treated exactly as a diploid outcross. These extra features, which are relatively simple add-ons to the core algorithms, are disabled by default and for clarity we do not present them in the present account. A brief summary is given in the supplement with more information in the Crosslink manual.

## Acknowledgements

We thank Eric van de Weg for helpful comments throughout the development of this package.

### Funding

This work was supported by the Biotechnology and Biological Sciences Research Council [BB/N006682/1, BB/M01200X/1 and BB/K017071/1].

*Conflict of Interest:* none declared.

